# Convpaint - Interactive pixel classification using pretrained neural networks

**DOI:** 10.1101/2024.09.12.610926

**Authors:** Lucien Hinderling, Guillaume Witz, Roman Schwob, Ana Stojiljković, Maciej Dobrzyński, Mykhailo Vladymyrov, Joël Frei, Benjamin Grädel, Agne Frismantiene, Olivier Pertz

## Abstract

We develop Convpaint, a universal computational framework for interactive pixel classification. Convpaint utilizes pretrained convolutional neural networks (CNNs) or vision transformers (ViTs) for feature extraction and enables easy segmentation across a wide variety of tasks. Available within the Python-based napari software ecosystem, Convpaint integrates seamlessly with other plugins into image processing pipelines, which we demonstrate with three workflows across different data modalities.

---

Many bioimage analysis pipelines start with a segmentation step. While deep learning methods (DL) offer high classification accuracy, they require extensive ground truth annotation data and dedicated hardware for training. Even foundation models, trained on more diverse data and expected to generalize to new applications without retraining, in practice still need retraining for basic research purposes [1–3]. In contrast, machine learning (ML) approaches using small models that can be trained interactively with sparse annotations, have proven to be highly effective (ilastik, Trainable Weka, Qupath, APOC) [4–7]. These approaches traditionally rely on hand-crafted filter banks to extract image features and train an ML model from sparse annotations and corresponding features, to predict the class of each pixel in the rest of the image or new images. While these models are quick to train and hand-crafted filter banks effectively describe texture or local image structures [8], breakthrough performance in capturing semantically meaningful information from images has been achieved through automatically learned filter banks, specifically convolutional filters in DL models [9]. Convpaint builds on this work, striking a balance between training speed, accuracy, and steerability by combining ML models that are fast to train with the power of DL. Instead of training a DL model from scratch, Convpaint extracts features from pretrained models like convolutional neural networks (CNN) or vision transformers (ViT) and uses them to train a random forest classifier. Convpaint’s design is modular, allowing the feature extractor to be easily replaced by any algorithm that returns local features from an input image (fig. 1A).

**Fig. 1.**
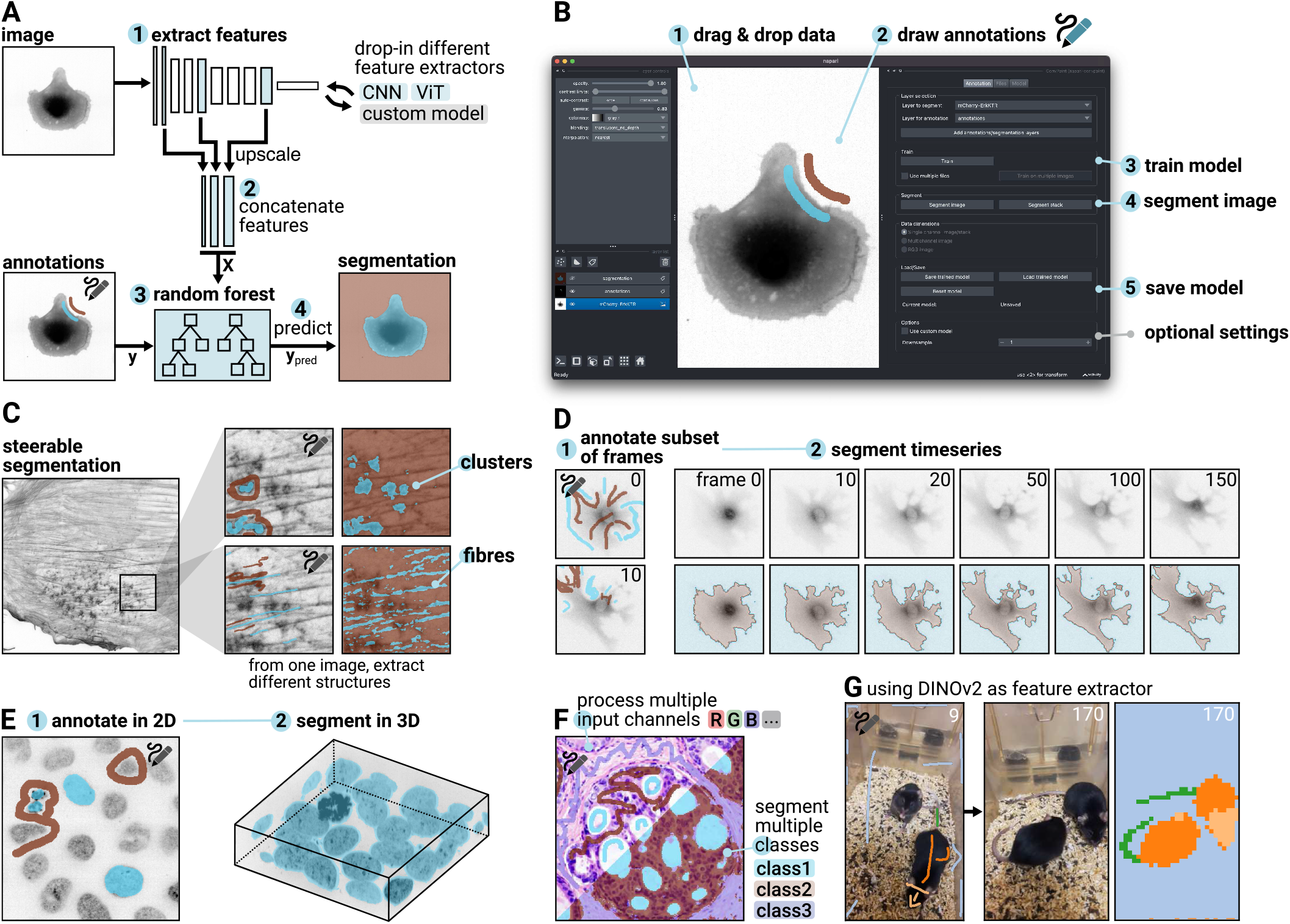
Overview of the Convpaint algorithm, user interface, and capabilities. **A** Convpaint architecture: 1) Features are extracted from multiple scalings of the input image using a pretrained neural network, 2) upscaled, and concatenated. 3) A random forest model, trained on sparse annotations, predicts the class for each pixel. Different feature extractor models can be used. **B** User interface in napari: 1) Supports various input formats. 2) Annotations drawn using the labels layer. 3) Single-click model training. 4) Single-click image segmentation with results displayed in a labels layer. 5) Model saving for future use. Advanced settings for custom models and normalization available in additional tabs. **C** Interactive adjustment of extracted structures based on annotations. **D** Segments time-series data across all frames with a single click, enabling immediate playback **E** Use napari’s visualization to verify 3D segmentation results. **F** Adapts to any number of input channels and output classes. **G** DINOv2 as a feature extractor allows the segmentation of macroscopic objects and scenes. Full data shown in fig. S5, suppl. movie M3.

Providing a graphical user interface within the napari ecosystem, Convpaint offers a straightforward way for researchers to repurpose pretrained DL models for their specific tasks without requiring coding or ML expertise (fig. 1B). Unlike neural networks trained to detect specific structures (e.g., spots, fibers), users can guide the Convpaint model by drawing sparse labels on regions of interest (fig. 1C). This interactive process, which can be completed in seconds, allows for iterative cycles of annotation and evaluation, rapidly improving the quality of segmentation results. Convpaint seamlessly handles multidimensional data, making it suitable for segmenting time series and 3D data (fig. 1D, E). It can be trained on an arbitrary number of input channels and output classes (fig. 1F, shown on synthetic data in S1). When coupled with different feature extraction models, Convpaint can be applied to bioimages across scales, from subcellular to cellular structures to animals (fig. 1G, S2). For experienced users, Python APIs are available, allowing them to programmatically control Convpaint. Convpaint incorporates several architectural optimizations that enhance training and prediction efficiency, setting it apart from similar software: it extracts crops around annotated pixels, minimizing unnecessary processing of entire images. It uses tiled parallel processing with appropriate padding for large images and stacks. Additionally, its integration with Dask allows for handling larger-than-memory files. The modular design makes customization easy, enabling users to integrate new feature extractors via simple functions, while Convpaint manages the user interface, classifier training, data management, and parallelization. The software is interoperable with a wide range of napari plugins, enabling complex image analysis workflows within a single software ecosystem without coding. We demonstrate this in three workflows.

**Workflow 1** highlights Convpaint’s capability to work with multichannel data. Imaging mass cytometry (IMC), spatial transcriptomics, or multiplexed immunofluorescence imaging, can image numerous biomarkers in the same sample. This provides a wealth of information posing new challenges for data analysis, particularly for interactive data exploration. Fig. 2A-D shows an exemplary use case on a 43-channel IMC dataset [10]. The data can be interactively loaded and browsed using napari-imc [11]. Instead of exporting the data for pixel classification in external software like ilastik as demonstrated in a previous study [12], pixel classification can be performed directly in napari using Convpaint. Here, we segment vein and surrounding tissue regions and identify markers that are differentially expressed between the two classes.

**Fig. 2.**
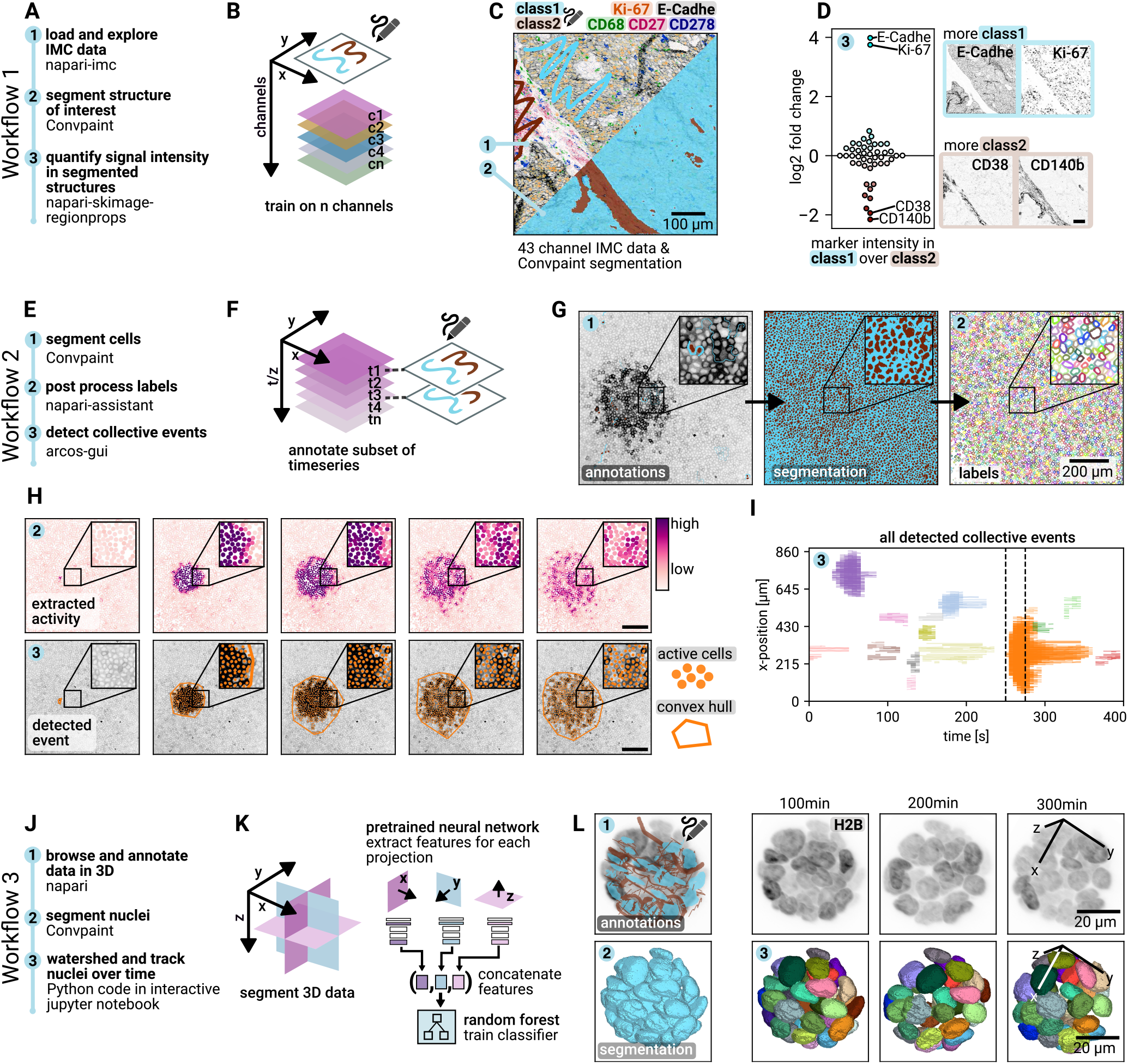
Image analysis workflows using Convpaint. Workflow 1 (multichannel dataset): **A** Example with multichannel IMC data. **B** Handles arbitrary input channels. **C** Interactive exploration with napari-imc. Labeled structures guide segmentation across all channels. Scale bar: 100 µm. **D** Use class labels for data exploration and statistical analysis, such as identifying differentially expressed markers. Scale bar: 100 µm. **Workflow 2 (time-series):** Supports 3D and time-series data. **E** Combined with arcos-gui to detect collective signaling events in MDCK cell movies. **F** Train classifier on multiple frames or z-slices to predict the rest. **G** Segmentation and post-processing with napari-assistant. Scale bar: 20 µm. **H** Example of collective event detection over 5 frames, with signaling activity and event overlay. Scale bar: 200 µm. **I** Overview of all detected events, with the period from panel H marked. **Workflow 3 (3D segmentation)**: **J** Segmentation of MCF10A acini from lightsheet microscopy, with 3D watershed instance segmentation and tracking with trackpy. **K** Feature extraction from multiple projections (xy, xz, yz) combined for random forest classification. **L** 3D rendering of tracked nuclei with color-coded IDs. Scale bar: 20 µm.

**Workflow 2** demonstrates Convpaint ability to analyze time lapse data with native support for interactive visualization in napari, to detect collective calcium signaling waves in an epithelial monolayer expressing a calcium biosensor (fig. 2E-I, suppl. movie M1) [13]. First, Convpaint is trained with scribbles on multiple frames to segment cells, then napari-assistant [14] is used for post-processing the labels and extracting biosensor information, and finally, ARCOS [15, 16] is utilized to detect and quantify collective signaling events.

To fully exploit information in 3D datasets using Convpaint, we implemented feature extraction from xy, xz, and yz projections. The features are concatenated to form the input for the random forest classifier. In **Workflow 3**, fig. 2J-L, this approach’s effectiveness is demonstrated on a lightsheet time lapse dataset of 3D mammary acini expressing a nuclear marker and an ERK biosensor. Following Convpaint segmentation, a simple 3D watershed is used for instance segmentation, and nuclei are tracked with an overlap-based algorithm (suppl. movie M2). Fig. S3 details how segmentation can be used to extract single-cell ERK activity signaling trajectories. Fig. S4 shows how information from multiple projections improves segmentation performance on a synthetic 3D dataset.

In this paper, we demonstrate that by using ViTs as feature extractors, which capture semantically richer information, we can accomplish tasks that were previously unattainable with traditional pixel classifiers. For example, leveraging DINOv2, pretrained on a large and diverse image dataset [17], we successfully segmented the head and tail of mice (fig. S5, suppl. movie M3) and detected whether their eyes were open or closed (fig. S6, M4). This approach is broadly applicable across animal species. With minimal annotations, we tracked body parts of sharks, even during complex movements such as rotations (fig. S7, M5).

Since there are no standard ground truth test datasets for evaluating interactive pixel classifiers, we developed a computational pipeline to generate test datasets for Convpaint. This pipeline automatically creates humanlike scribbles from existing segmentation dataset annotations, allowing us to quantitatively assess Convpaint’s segmentation performance and compare different feature extractors. It also enables testing Convpaint across varying levels of annotation coverage. We observe significant improvements in segmentation performance on complex data, such as detecting cancerous tissue in histology slides, when using pretrained neural networks as feature extractors. The pipeline used to generate the test data, the results across different datasets, and insights into how performance depends on the number of scribbles are discussed in detail in the supplementary information and shown in fig. S8. Randomly sampled results for all datasets are shown in figs. S9, S10, S11.

The results show Convpaint’s flexibility and performance, which, together with its seamless integration within the napari ecosystem, makes it an attractive segmentation tool for a wide variety of image analysis tasks and data types. We envision Convpaint empowering researchers across diverse fields, from cell biologists to ecologists. With no programming knowledge required, users can access state-of-the-art, steerable segmentation tailored to their specific needs with just a few clicks. This accessibility brings advanced image analysis techniques to a broader scientific community.

The code is open source (BSD-3), available on GitHub^1^, and can be installed from the napari hub^2^ or PyPI^3^. It runs on all common operating systems and standard consumer hardware, with optional GPU acceleration. We provide installation instructions, documentation, and video tutorials^4^.

## Supporting information

Movie M1: Detection of calcium waves in MCDK cells

Movie M2: Segmentation of MCF10A acini

Movie M3: Segmentation of mouse body parts

Movie M4: Detection of open/closed eyes in humans and mice

Movie M5: Segmentation of shark body parts

## ACKNOWLEDGEMENTS

This work has been supported by the Chan Zuckerberg Initiative (CZI) grant NP2-0000000095 to LH and OP, Uniscientia fellowship 187-2021 to OP, and Schweizerischer Nationalfonds (SNF) grant 310030_185376 to OP. We thank the Scientific Center for Optical and Electron Microscopy (ScopeM) of ETH Zurich, Switzerland, for access to their instruments and services and Dr. Tobias Schwartz for his assistance in acquiring lightsheet data. Calcium imaging data was kindly provided by Yasuto Takeuchi and Yasuyuki Fujita. Other microscopy experiments were performed on equipment supported by the Microscopy Imaging Center (MIC), University of Bern, Switzerland. The mouse icon in A by DBCLS https://togotv.dbcls.jp/en/pics.html is CC-BY 4.0 licensed.

## AUTHOR CONTRIBUTIONS

LH conceptualized the work. LH, GW, RS, AS, MD, MV, and BG contributed to the development of the software and documentation. RS quantified performance. JF, LH, BG, RS, and AF acquired data. Figures were created by LH. LH and OP wrote the manuscript and acquired funding. All authors read and approved the final manuscript.

## Methods

### Convpaint implementation details

Convpaint features a modular architecture designed to accommodate a wide range of feature extractors, enhancing existing algorithms or pretrained models with added steerability. We compare three different types of feature extractors:

### CNN

We utilize the VGG16 [18] architecture implemented in pytorch [19], pretrained on the ImageNet dataset, to extract local image features such as edges, textures, and color channel correlations when working with RGB images. Downscaled versions of the input image are passed through VGG16, creating a featurized image pyramid. These features are then upscaled and concatenated with unscaled outputs and deeper CNN layer features, balancing segmentation speed and accuracy. This method effectively generalizes to a variety of image segmentation tasks (fig. S2). We evaluated different configurations of input scalings and layers for feature extraction and provide default settings that perform well on all of the tested datasets.

### ViT

We incorporate two ViT models — DINOv2 [17] and UNI [2]. DINOv2 is pretrained on 142M images from ImageNet, while UNI is pretrained on a large histology dataset. These models extract patch features of 14×14 pixels (DINOv2) and 16×16 pixels (UNI), providing superior performance in certain segmentation tasks despite a loss in resolution for fine details below the patch size, like small cell protrusions. For each patch, the ViT-S/14 distilled DINOv2 model we used extracts 384 features, while UNI extracts 1024 features. For all DINOv2 experiments, we utilized the variant with registers, as this configuration produced less patch noise in the predictions (fig. S8J).

### Classical Filter bank

To compare the performance of Convpaint when using pretrained neural networks versus classical filter banks as feature extractors, we employed the filters implemented in napari-ilastik^5^. We chose the maximal combination of filters and sigma parameters suggested in the library, including Gaussian, Laplacian of Gaussian, Gaussian gradient magnitude, difference of Gaussians, structure tensor eigenvalues, and Hessian of Gaussian eigenvalues, with sigma values 0.3, 0.7, 1.0, 1.6, 3.5, 5.0, 10.0. Fig. S12 shows a visual comparison of filters used in classical filter banks versus learned convolutional filters extracted from VGG16.

Convpaint is optimized for both training and prediction efficiency:

- Crops Around Annotations: Avoids processing entire images by extracting crops around annotated pixels.
- Tiling and Parallel Processing: Handles large images by tiling them and using parallel processing, with appropriate padding to minimize edge effects. One-click batch processing for image stacks.
- Data Management: Manages larger-than-memory files using Dask, appropriate handling of additional image dimensions (channels vs. time/z-slices)
- Customizability: Users can easily integrate other feature extractors by implementing a simple function that returns a feature matrix from an image. Convpaint takes care of the user interface, classifier training, data management, and parallelization.

For users wanting to implement a custom feature extractor, we provide a blueprint as a starting point. Convpaint makes it easy to take existing architectures and repurpose them in minutes. To give another example, we have explored using intermediate outputs of a cellpose model as a feature extractor. While the model is trained to predict cell masks, in Convpaint it can be steered to segment cell boundaries or nuclei with a couple of scribbles. Another possibility is to simply concatenate the output of multiple feature extractors, which allows combining their strengths. As an example, we successfully combined the pixel-accurate segmentation of VGG16 with the semantic understanding of DINOv2 S13: while DINOv2 reliably differentiates shark body parts, it is limited in accuracy by the patch size of its features. VGG16, on the other hand, precisely masks the shark but lacks the semantic understanding to correctly label different anatomical structures. Combining both models achieves semantically correct labels with high spatial resolution.

The design of Convpaint streamlines experimentation with different feature extractors and ensures that new innovations in computer vision can be easily integrated to leverage its performance.

### Quantification of segmentation performance

Assessing Convpaint’s performance, especially given its interactive nature, is challenging. Even non-interactive models face problems in unbiased performance evaluation in bioimage analysis [20]. Given a lack of scribble-annotated datasets, we created an algorithm to generate human-like scribbles from existing ground truth datasets, allowing for an unbiased quantitative assessment of segmentation performance. For each image from the data set, we generated scribble masks with varying annotation densities (fig. S8A). We evaluated three datasets: cellpose [21], foodseg103 [22], and a subset of a breast cancer histology slide database [23]. We chose the foodseg103 dataset based on the hypothesis that classical filters would struggle to assign semantic information for items containing highly variable textures. Similarly, we selected the breast cancer dataset, representing a common challenging use case in biological research. In total, we evaluated segmentation performance on 148k samples. The code to automatically generate scribbles and recreate the figures is available on GitHub^6^. The repository also contains the full results, including multiple performance metrics for each image and classifier at different levels of scribble annotations, as well as plots exploring the effects of feature extractor parameters on segmentation.

#### Scribble generation

To closely mimic human annotations, scribbles are created by combining three types of algorithmically generated lines:

1. Center ridge lines: Sampled from the primary skeleton of the ground truth mask.
2. Boundary parallel lines: Sampled from the secondary skeleton, which is derived from the ground truth mask after subtracting the primary skeleton.
3. Boundary perpendicular lines: Lines connecting the primary skeleton to the mask boundary.

By varying the sampling density, we can generate different levels of annotation coverage, such as 0.1% or 1% of the image pixels. The algorithm can also vary scribble type, length, and width, making it versatile for research scenarios beyond the scope of this study, e.g. how scribble types affect segmentation performance. For the cellpose dataset, which consists mostly of images with numerous small, cell-like objects, we generated a large number of short, 1-pixel-wide scribbles. The ground truth masks were converted from instance segmentation to semantic segmentation (i.e., foreground/background instead of cell IDs). For the foodseg103 dataset, which features fewer but larger regions of different food items, we generated fewer, longer scribbles with a width of 2 pixels. For the histology dataset, we created medium-length scribbles with 2 pixels width.

#### Dataset evaluation

As quantitative readout for segmentation performance we report the mean intersection over union (mIoU). For the cellpose dataset (540 images, fig. S8B), which involves segmenting small cell-like structures from a dark background, we observe similar performance between VGG16 filters and classical filter banks (fig. S8C). Increased performance variability in low-annotation regimes can be attributed to the random information content of the few annotated pixels. At higher annotation levels, mIOU is often limited by imprecisions in the ground truth annotations, such as missing protrusions (fig. S8D). DINOv2 performs poorly due to its patch size being too large for resolving smaller cellular details of a few pixels. Randomly selected sample images and predictions are shown in fig. S9.

To reduce computation time for the foodseg103 dataset (fig. S8E) with 4983 images, we excluded images larger than 640k pixels and used 520 images sampled from the dataset for evaluation. Segmentation of the images requires distinguishing food items with complex textures and colors. VGG16 filters outperform classical filter banks for all tested configurations. When only providing very sparse annotations, smaller networks with VGG16 filters show better performance, likely due to reduced overfitting. Because of the larger label regions, unlike in the cellpose dataset, DINOv2’s performance is much less constrained by patch size and the ViT massively outperforms both classical and VGG16 filters, even for very low annotation regimes (fig. S8F). For all models, we observe diminishing returns in segmentation performance as annotation levels increase. For Convpaint users, this suggests that adding a few more annotations is beneficial when only a minimal number of scribbles are present, but if performance gains plateau, it’s a good time to stop (shown for a VGG16 model in fig. S8G). Randomly selected sample images and predictions are presented in fig. S10.

Motivated by DINOv2’s performance on foodseg103, we tested it on a subset of a breast cancer histology slide dataset (fig. S8H). DINOv2 again clearly outperforms both classical filter banks and VGG16 filters. We also evaluated a histology-specific model built with the same architecture as DINOv2 [2], which surprisingly shows no advantage over DINOv2 (fig. S8I). Predictions from different models (8 out of 10 images tested) are shown in fig. S11. DINOv2 achieves the highest mIoU results and produces visually less noisy predictions compared to other models.

In summary, DINOv2 demonstrates superior performance in segmentation tasks that require broad contextual information when patch-sized resolution is sufficient, generalizing well across domains. In contrast, VGG16 and classical filters provide precise segmentation when local information is adequate for the given task. Here, VGG16 generally matches or exceeds the performance of classical filters, making it a suitable alternative.

## Methods and data availability

### Workflow 1: IMC multichannel data

IMC data from [10], available on Zenodo^7^ were loaded using the napari-imc plugin [11]. Convpaint was trained on one FOV (fig. 2 shows Patient 01, Panorama 02, Position 1-1). Skimage was used to extract the per-channel statistics for the segmented regions. The log2-fold change in signal intensity was calculated using NumPy and plotted with Matplotlib. Step-by-step instructions for recreating the workflow are available in the documentation.

### Workflow 2: MDCK calcium waves

Time lapse data sets of calcium signaling waves were obtained from MDCK epithelial cells that stably express GCaMP6S - a genetically encoded intracellular calcium sensor (imaging data courtesy of Yasuyuki Fujita). The movies were loaded into napari and segmented using Convpaint. The resulting binary masks were processed with ARCOS [15] to detect and quantify collective signaling events. The code to recreate the figures is available on GitHub TODO. We have made the raw imaging data available on the BioImageArchive [24] under the accession number S-BIAD1135. Step-by-step instructions for recreating the workflow are available in the documentation.

### Workflow 3: 3D segmentation and nuclei tracking

We demonstrate that Convpaint can effectively segment and track single cells within a dense 3D spheroid. In fig. S3, we illustrate how this data can be further processed to extract single-cell ERK signaling activity dynamics. The cells used are MCF10A, expressing a histone H2B nuclear marker and ERK-KTR, which reports ERK activity through reversible nucleus/cytosol translocation following phosphorylation by active ERK [25] (scheme in fig. S3A). Data were acquired using a lightsheet microscope with a 5-minute resolution and an isotropic voxel size of 0.145 µm. Raw imaging data and protocols are available on BioImageArchive [26] under accession number S-BIAD1134.

### Other figures

The 3D nuclear data in fig. 1F is part of the scikit-image [27] data module, called *cells3d*, originally provided by the Allen Institute for Cell Science. The synthetic data in fig. S4 was generated by another group using FiloGen [28] and is available from the Broad Bioimage Benchmark Collection (BBBC046^8^) [29], showing cell PD-ID451/AR1/T024.

The datasets used for performance quantification have all been previously published. Fig. S8B shows the cellpose dataset [21]; fig. S8C shows the foodseg103 dataset [22]; and fig. S11 uses data from the Breast Cancer Semantic Segmentation (BCSS) dataset^9^ [23]. The movie shown in fig. S5 is a supplement^10^ to a study on epileptic behavior in mice [30]. Movies in figs. S6A, B and S7 were acquired by the authors and are available upon request. The movie in fig. S6D, E is available online^11^ for educational purposes under the Mixkit Restricted License. Images in fig. S2 were acquired by the authors and are available upon request, except for the histology slide images, which are from Wikimedia Commons^12^, or provided by the scikit-image library, acquired at the Center for Microscopy And Molecular Imaging (CMMI).

## Supplementary figures

- S1: Correlation extraction across multiple color channels.
- S2: Image segmentation across diverse domains.
- S3: Measuring ERK signaling dynamics at the single-cell level in MCF10A acini.
- S4: Improved segmentation performance using multiple projections.
- S5: Detecting mouse body parts in video.
- S6: Eye state detection in humans and mice.
- S7: Detecting shark body parts in video.
- S8: Quantification of segmentation performance and model comparison.
- S9: Feature extractor performance on the cellpose dataset.
- S10: Feature extractor performance on the foodseg103 dataset.
- S11: Feature extractor performance on histology slides.
- S12: Visual comparison of handcrafted vs. learned filters.
- S13: Combining VGG16 and DINOv2 features for enhanced spatial precision at mask boundaries while maintaining semantic information.

## Supplementary movies

- M1: Detection of calcium waves in MCDK cells
- M2: Segmentation of MCF10A acini
- M3: Segmentation of mouse body parts
- M4: Detection of open/closed eyes in humans and mice
- M5: Segmentation of shark body parts

**Fig. S1.**
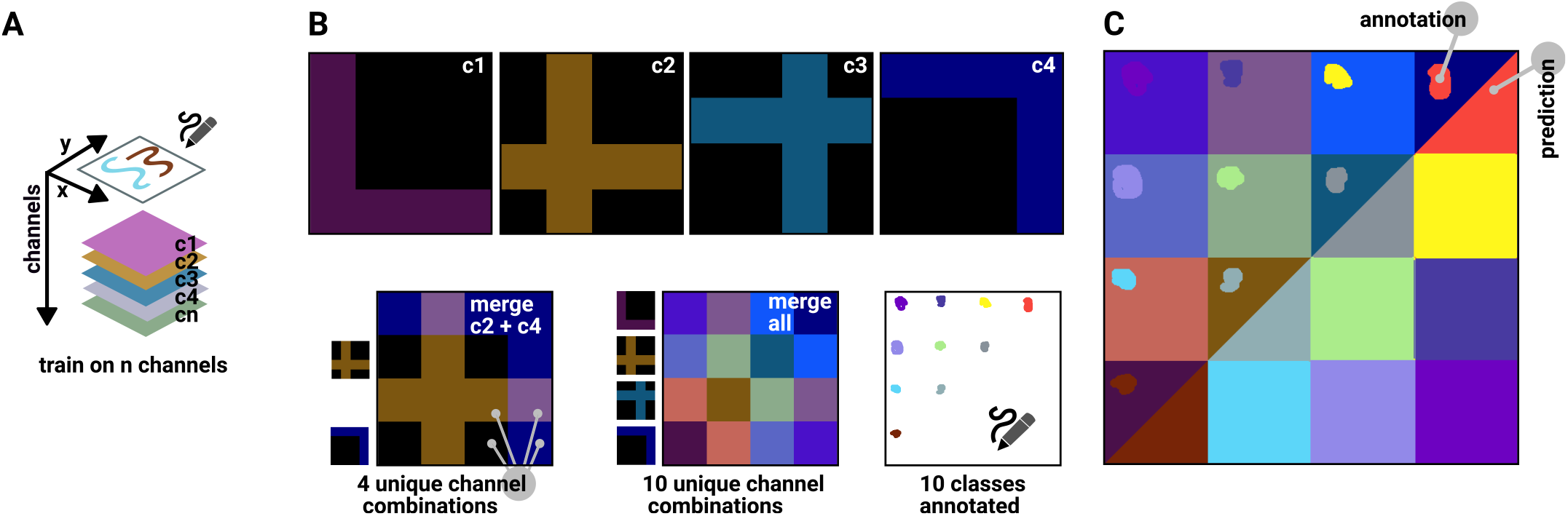
Correlation extraction across multiple color channels. **A** The Convpaint classification algorithm can process an arbitrary number of input color channels. **B** This is demonstrated on an artificial image with four channels, which when merged lead to 10 unique color combinations. These 10 combinations are labeled with 10 class labels that can only be reconstructed if the algorithm takes into account the interplay of the different channels. **C** Convpaint correctly predicts the correct class label for pixels that were not labeled, with minor artefacts on boundaries between squares.

**Fig. S2.**
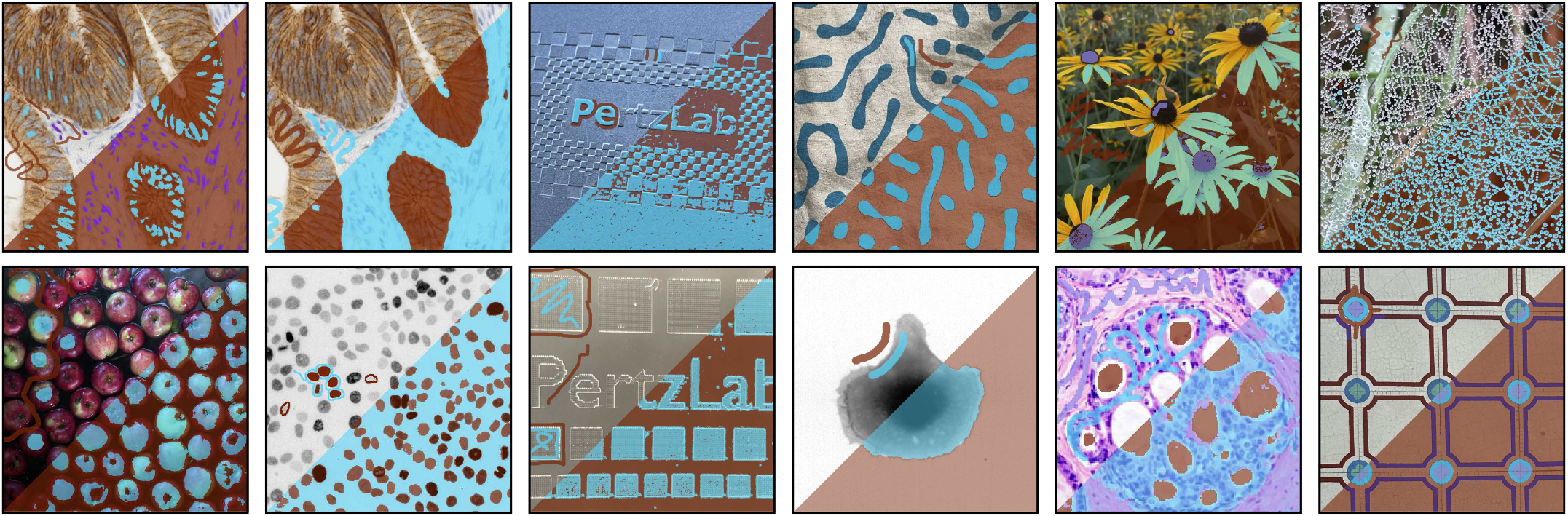
Image segmentation across diverse domains. All images use VGG16 with the default configuration as feature extractor.

**Fig. S3.**
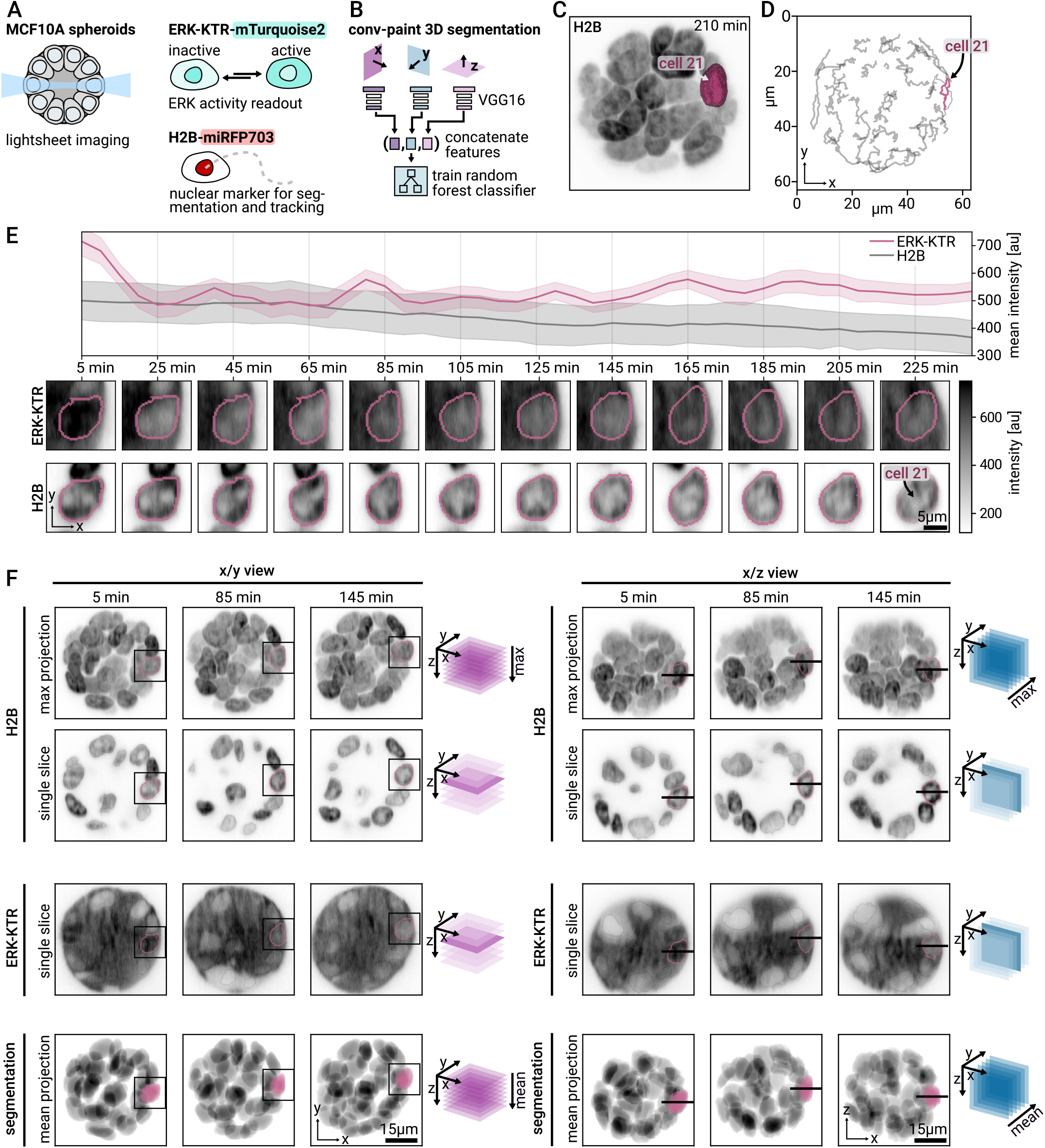
Measuring ERK signaling dynamics at the single-cell level in MCF10A acini. **A** Spheroids are imaged with a lightsheet microscope. The cells express an ERK activity sensor and nuclear marker for segmentation and tracking. **B** Convpaint is used to segment the nuclei in 3D. **C** Panels C-F track a single cell in the spheroid over time, here its mask is shown overlaid on a 3D max projection. **D** Tracks of all cells from 0 to 250 minutes, selected cell highlighted in color. **E** Mean nuclear ERK-KTR intensity over time as a proxy for ERK activity. We see ERK trajectories as previously described by Ender et al. [25]. In comparison, the mean intensity of the nuclear marker shows some bleaching but no fluctuations otherwise. The images show crops around the selected cell (mean of 3 z-slices, [+1,0,-1] around the z position of the cell centroid). Scale bar is 5 µm. **F** Highlighting the tracked position of the cell within the spheroid for different time points, projections, and channels. Box shows insets in panel E. Scale bar is 15 µm.

**Fig. S4.**
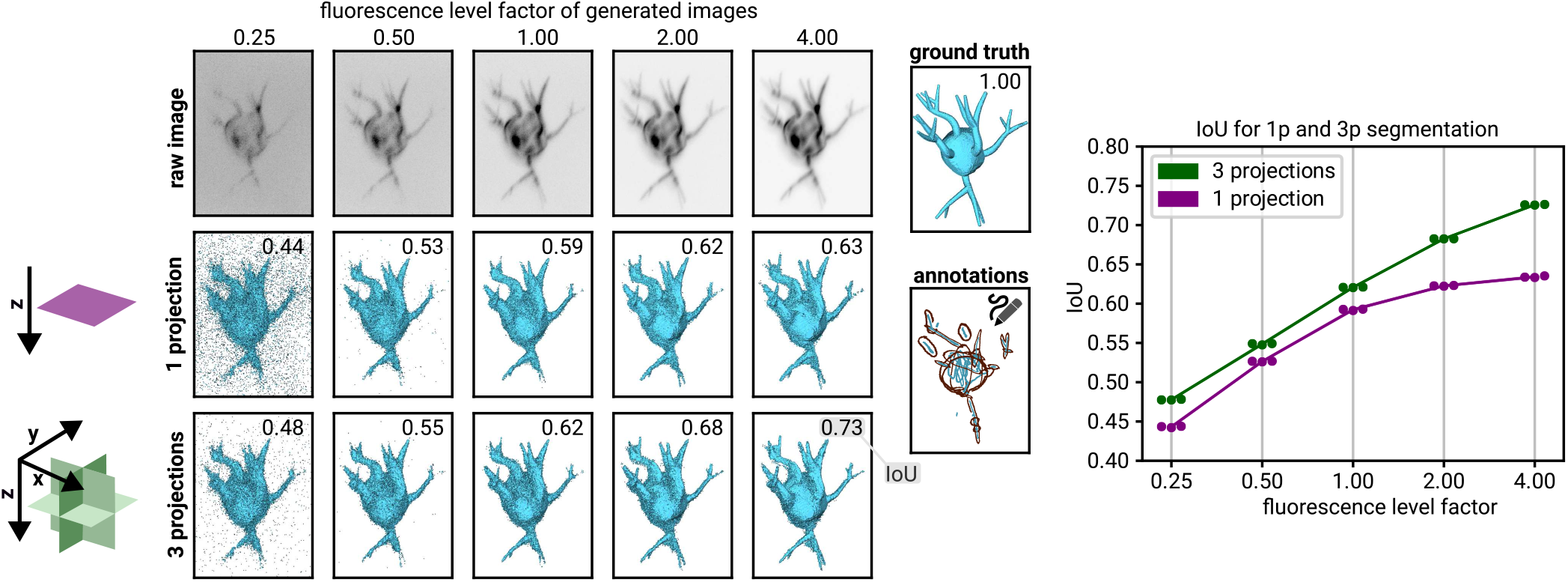
Improved segmentation performance using multiple projections. Convpaint segmentation performance compared on an artificial cell when extracting features from 1 projection (purple) versus concatenating 3 projections (green), using VGG16 with default configuration as feature extractor. Different signal-to-noise regimes are tested, which are configured by the fluorescence level factor (0.25-4) in the FiloGen software. Performance is measured as intersection over union (IoU). Using 3 projections leads to better segmentation results for all fluorescence level factors. A larger increase in performance is observed for images with a better signal-to-noise ratio.

**Fig. S5.**
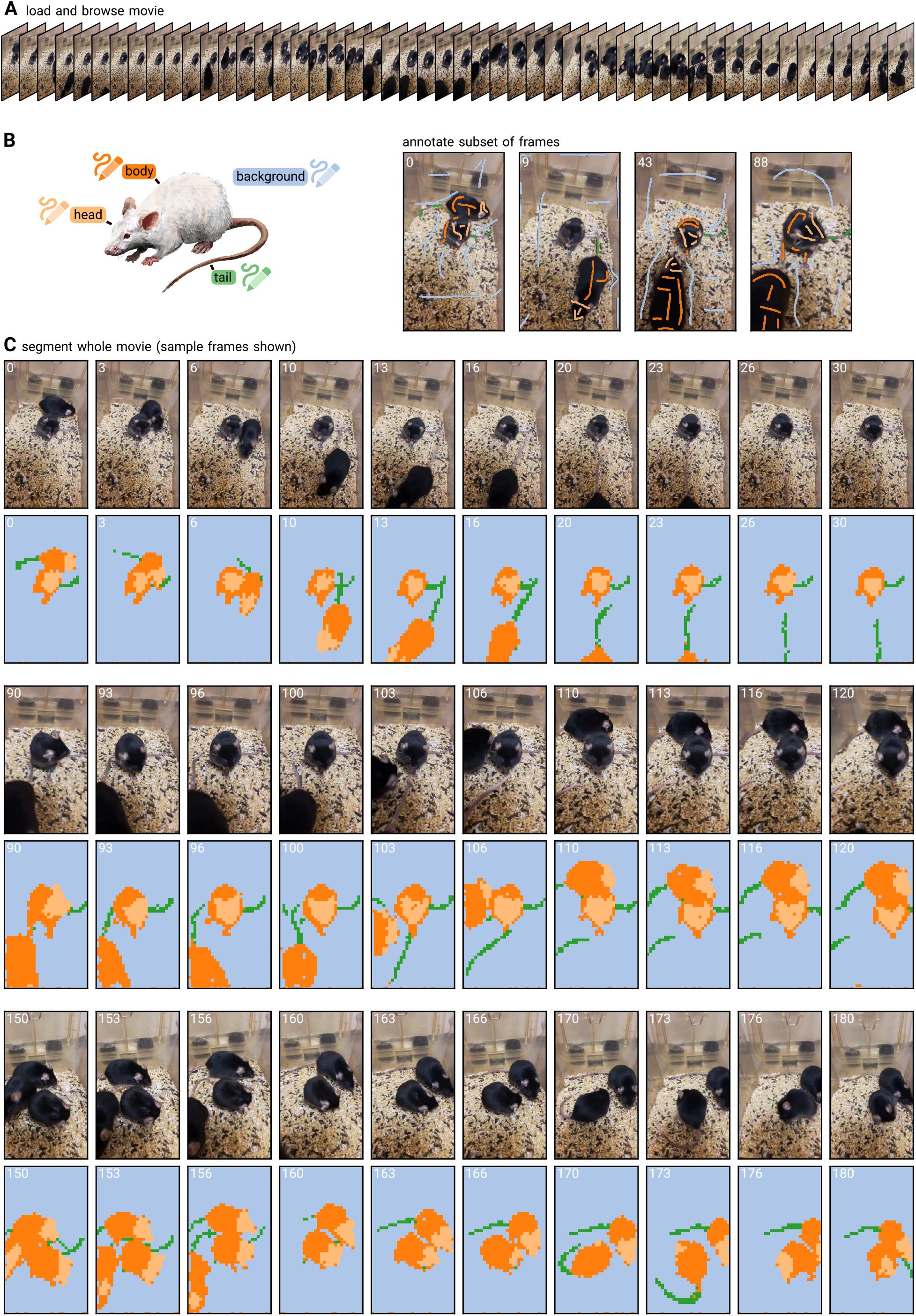
Detecting mouse body parts in video. Corresponding movie shown in suppl. M3. **A** The movie shows mice moving in a cage. The camera is handheld and the mice move in and out of frame. Some frames contain motion blur. **B** Convpaint with DINOv2 as feature extractor is trained on scribbles of four different frames. Head, body, tail, and background are annotated. **C** The trained model is used to predict the rest of the movie.

**Fig. S6.**
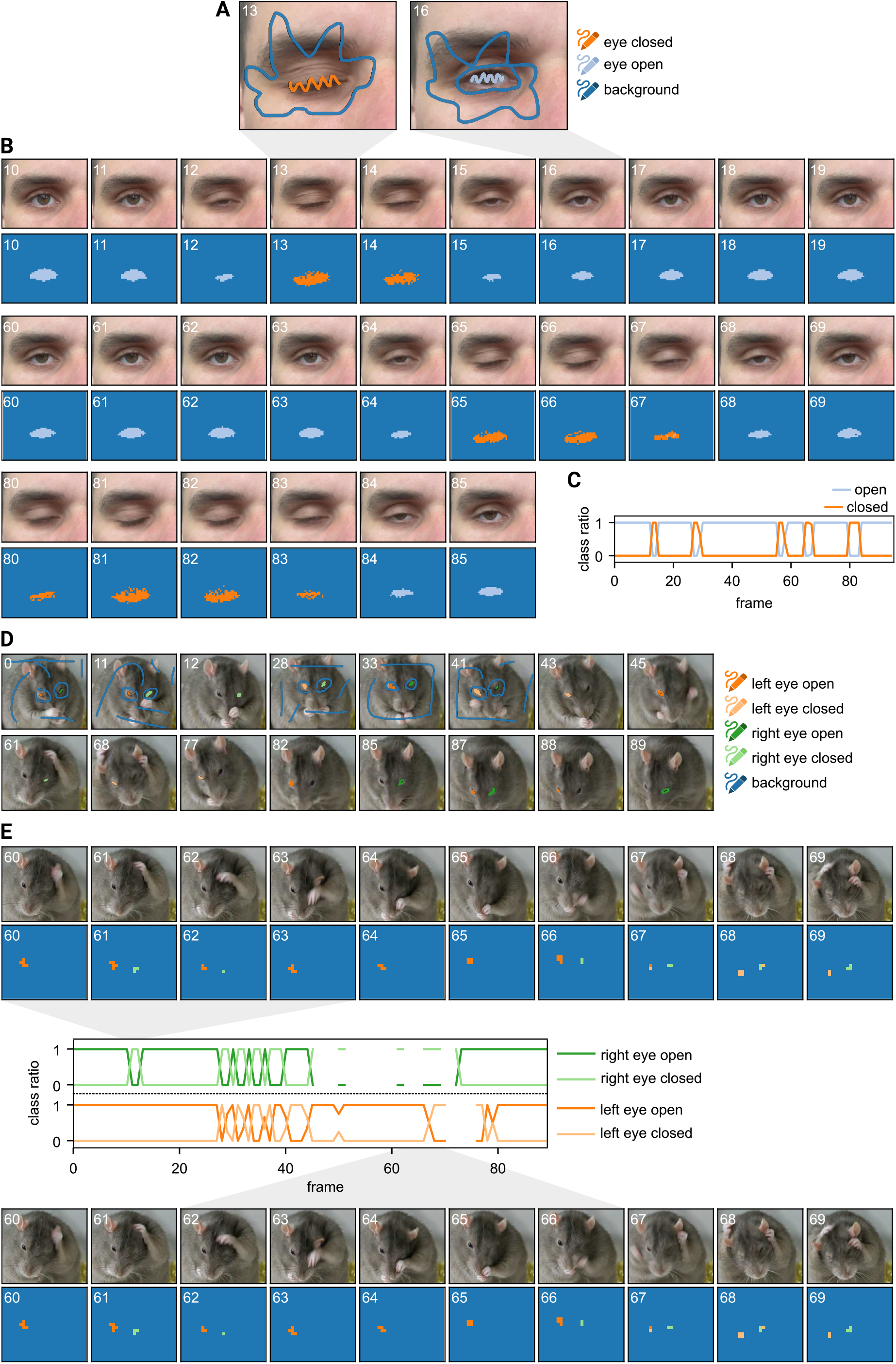
Eye state detection in humans and mice. Corresponding movie shown in suppl. M5. **A** Two frames are labeled with three classes (eye open, eye closed, background) **B** Convpaint is used to segment the rest of the movie. Sample frames are shown. **C** Quantification of class abundance. **D** Same as A but for a mouse. The left / right eyes are classified separately. **E** Sample frames and quantification of segmentation. No data is shown if no pixels are detected (mouse is grooming and a limb is obstructing the eyes).

**Fig. S7.**
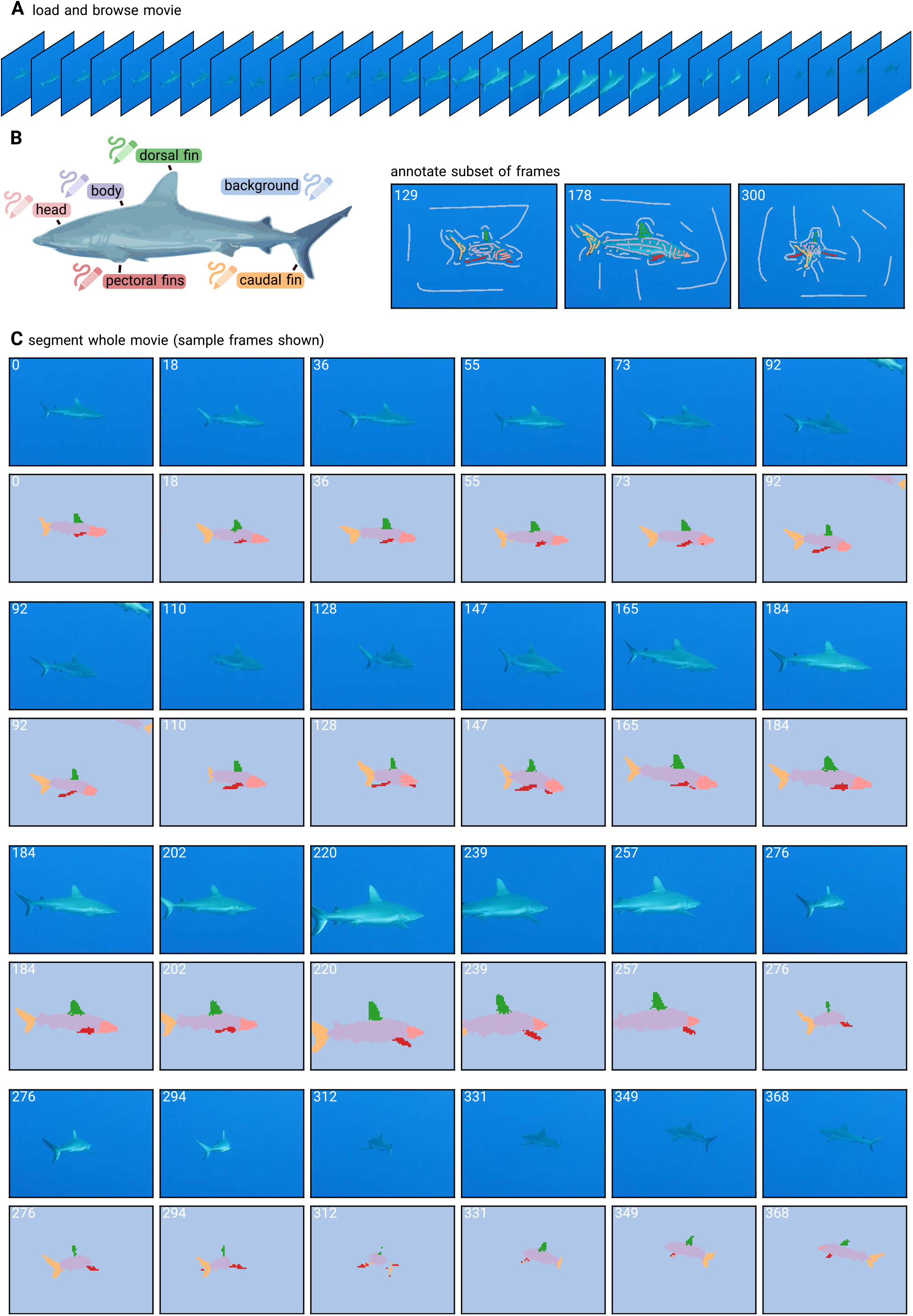
Detecting shark body parts in video. Corresponding movie shown in suppl. M4. **A** The movie shows a shark swimming from multiple angles. Camera is hand-held. **B** Convpaint with DINOv2 as feature extractor is trained on scribbles on three different frames. Head, body, dorsal, caudal, pectoral fins, and background are annotated. **C** The trained model is used to predict the rest of the movie.

**Fig. S8.**
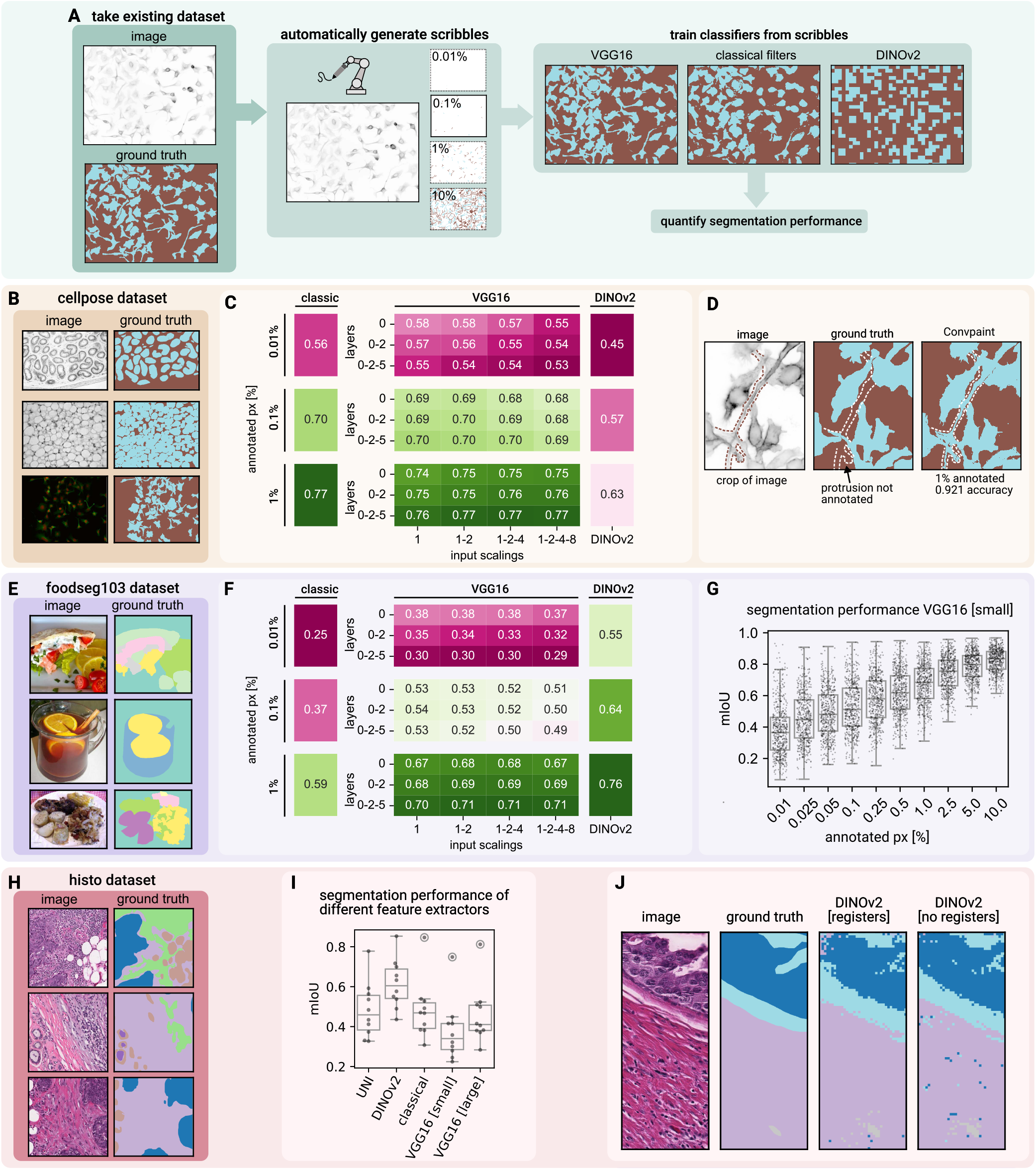
Quantification of segmentation performance and model comparison. **A** Ground truth datasets are used to automatically generate scribbles to train Convpaint. The results are then evaluated against the ground truth to quantify the performance of different feature extractors. Various feature extractors are tested, including VGG16 layers with different input scalings, classical filter banks with all napari-ilastik filter/sigma combinations, and DINOv2 ViT-S/14. **B** Testing on the cellpose dataset (three sample images and masks shown). See fig. S9 for segmentation results. **C** Mean mIoU scores for different annotation levels in the cellpose dataset. Similar performance for classical filters and VGG16. DINOv2 underperforms due to large patch size limitations in capturing small cellular details. **D** Model performance quantification scores can be limited by the quality of ground-truth annotations. A cell protrusion, missing in the ground truth, is correctly segmented by the model. **E** Testing on the foodseg103 dataset (three samples shown). See fig. S10 for segmentation results. **F** Mean mIoU scores for different annotation levels on the foodseg103 dataset. Both pretrained architechtures (CNN, ViT) outperform classical filter banks as feature extractors across all annotation regimes. DINOv2 has highest mIoU scores. In this dataset, the model’s performance is less affected by patch size because of larger regions of interest. **G** For all models, diminishing returns on segmentation performance can be observed with increasing annotation levels, here shown for VGG16 (layer 0 and input scalings 1,2) on the foodseg103 dataset. **H** Testing on a histology slide dataset. See fig. S11 for segmentation results. **I** Mean mIoU scores for different annotation levels in the histology slide dataset. DINOv2 outperforms all other models, including the histology-specific model UNI. **J** DINOv2 with registers produces predictions with less patch noise (not quantified).

**Fig. S9.**
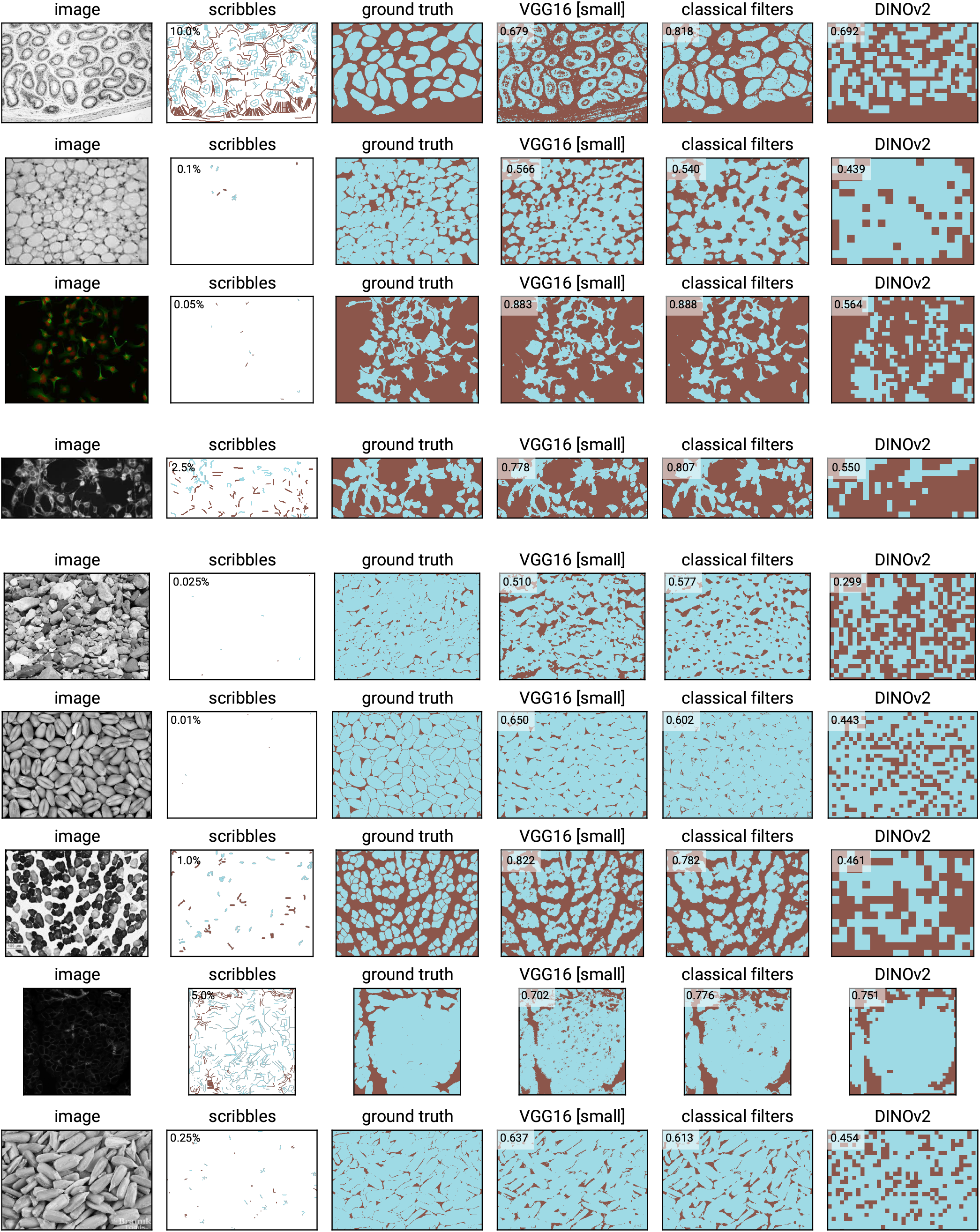
Feature extractor performance on the cellpose dataset. Randomly selected images. For plotting, the scribbles were dilated for better visibility. The number in the upper left corner of the prediction images shows the mIoU score.

**Fig. S10.**
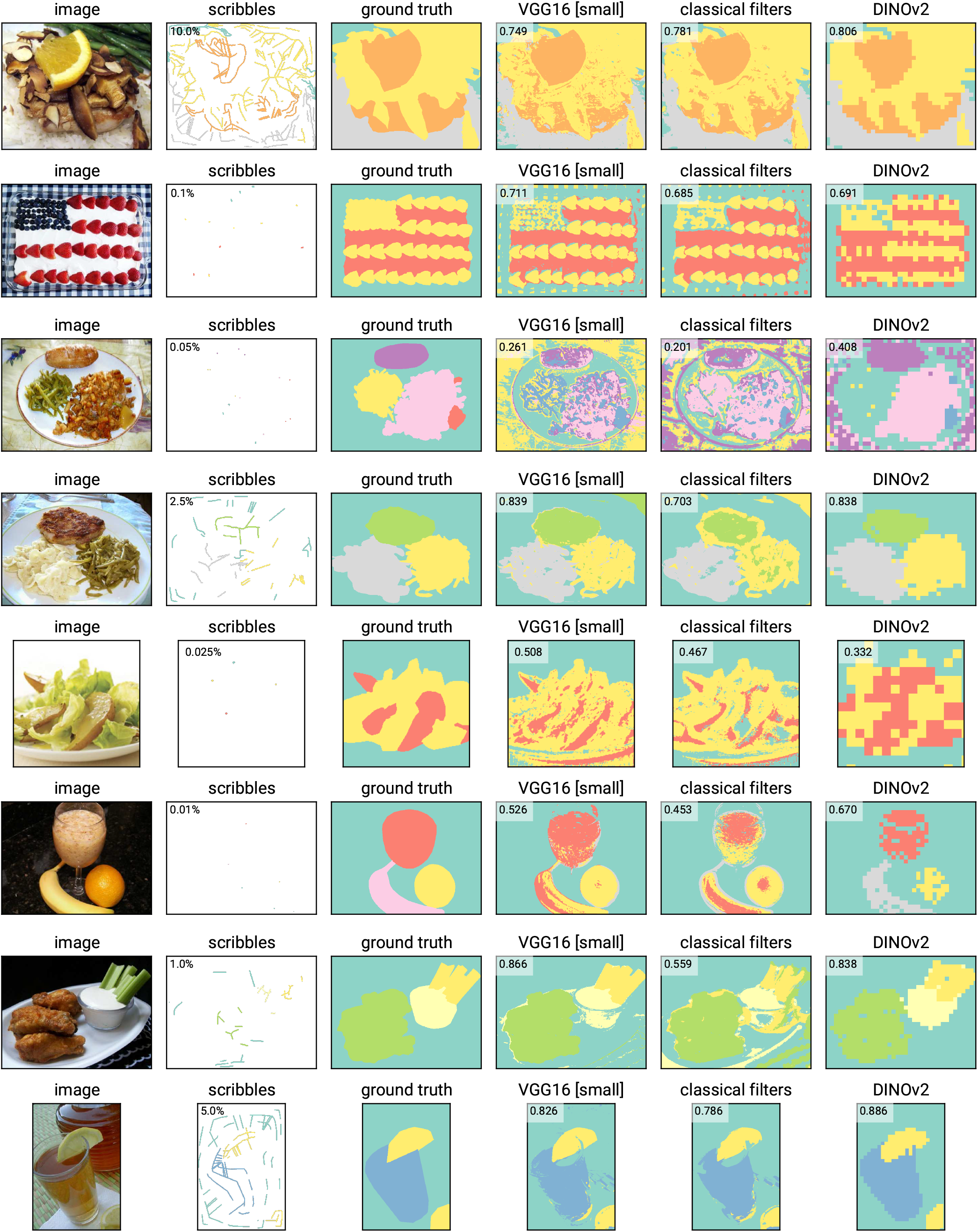
Feature extractor performance on the foodseg103 dataset. Randomly selected images. For plotting, the scribbles were dilated for better visibility. The number in the upper left corner of the prediction images shows the mIoU score.

**Fig. S11.**
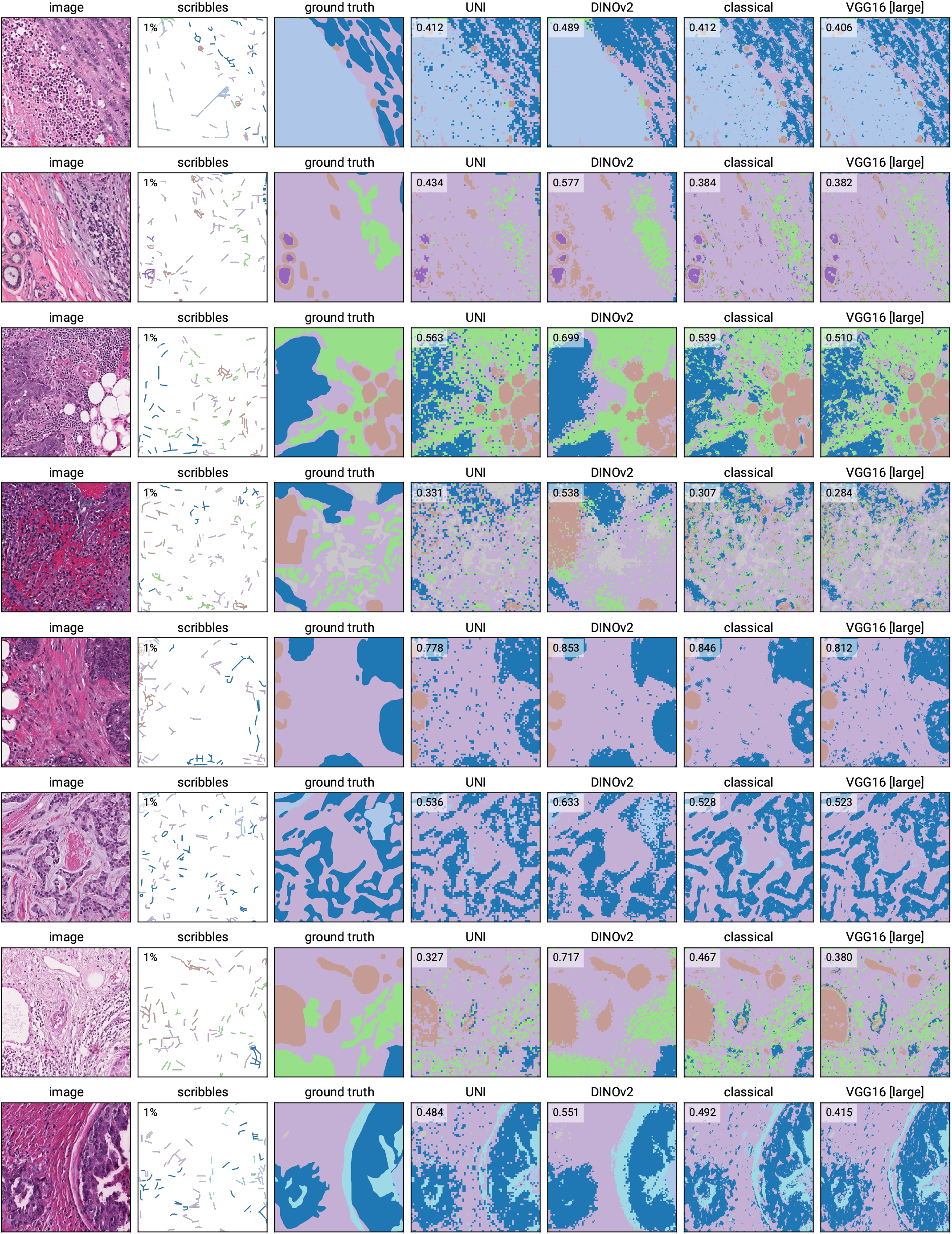
Feature extractor performance on histology slides. Scribbles were automatically generated from expert annotated ground truth. For plotting, the scribbles were dilated for better visibility. Eight out of ten images used for evaluation in fig. S8I are shown. The number in the upper left corner of the prediction images shows the mIoU score. DINOv2 outperforms all other models in this dataset.

**Fig. S12.**
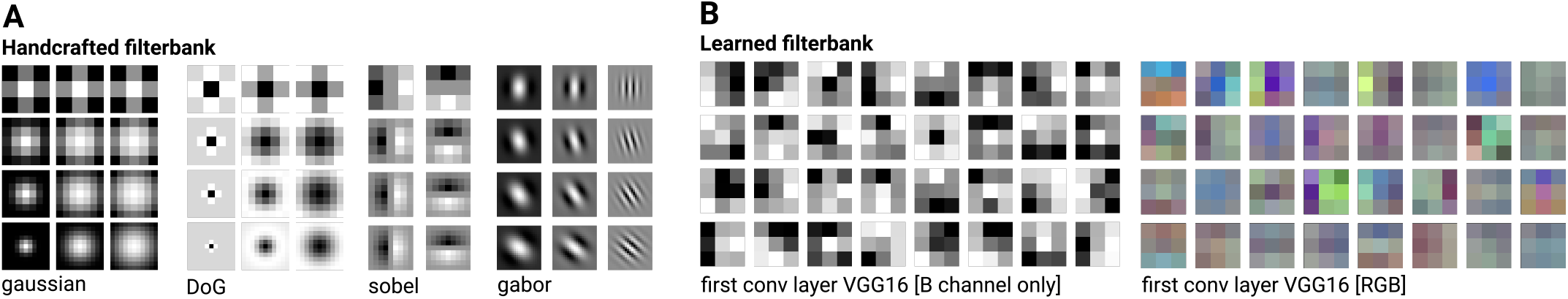
Visual comparison of handcrafted vs. learned filters. **A** Filters used classically in handcrafted filter banks. Here we show examples of filter kernels with different parameters for Gaussians, Difference of Gaussians (DoG), Sobel, and Gabor. While these handcrafted filter banks are more interpretable, the patterns they extract often overlap, leading to redundancy among the filters. **B** Filters extracted from the first convolution layer of a CNN (VGG16) network. The filters have a 3×3×3 shape, which makes them intrinsically capable of extracting correlations between color channels in RGB images. Although VGG16 filters are less interpretable, they are computationally optimized to extract orthogonal image features that are useful for image classification.

**Fig. S13.**
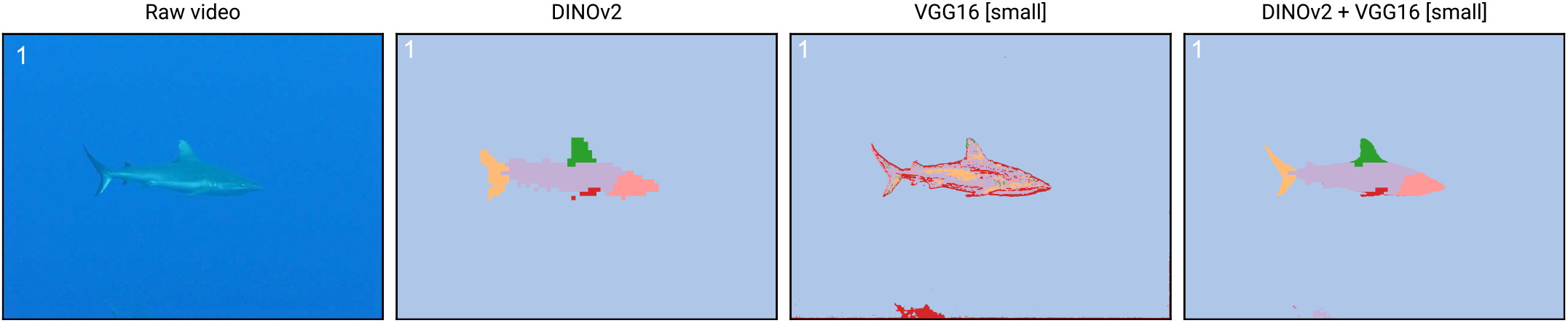
Combining VGG16 and DINOv2 features for enhanced spatial precision at mask boundaries while maintaining semantic information. While DINOv2 features excel at capturing abstract semantic information at the patch level, VGG16 features are good at capturing local spatial information at the pixel level. By concatenating the features of both models, we can leverage the strengths of both models.

## Footnotes

1 www.github.com/guiwitz/napari-convpaint

2 https://www.napari-hub.org/plugins/napari-Convpaint

3 https://pypi.org/project/napari-convpaint/

4 https://guiwitz.github.io/napari-Convpaint/book/Landing.html

5 https://github.com/ilastik/ilastik-napari

6 https://github.com/quasar1357/scribbles_creator

7 https://doi.org/10.5281/zenodo.5555575

8 https://bbbc.broadinstitute.org/BBBC046

9 https://bcsegmentation.grand-challenge.org/BCSS/

10 https://doi.org/10.3389/fgene.2018.00581.s007

11 https://mixkit.co/free-stock-video/gray-and-white-rat-32060/

12 https://commons.wikimedia.org/wiki/File:Breast_DCIS_histopathology_(1).jpg

